## Supplemental material for "First-in-human evaluation of cutaneous innate and adaptive immunomodulation by mosquito bites"

**SUPPLEMENTARY INFORMATION**

### Supplementary Tables

| Characteristic | Total | Immunohistochemistry | RNASeq | Flow Cytometry |
|---|---|---|---|---|
| n | 30 | 10 | 10 | 10 |
| Age, median years [IQR] | 32.5 [21.75, 37] | 33.5 [22.0, 35.0] | 33.5 [25.5, 38.5] | 30.0 [21.0, 36.0] |
| Female | 13 (43.3) | 5 (50.0) | 5 (50.0) | 3 (30.0) |
| Occupation | | | | |
| Businessman/seller/ entrepreneur | 3 (10.0) | 0 (0.0) | 0 (0.0) | 3 (30.0) |
| Government Employee | 1 (3.3) | 0 (0.0) | 0 (0.0) | 1 (10.0) |
| Factory Worker | 1 (3.3) | 0 (0.0) | 1 (10.0) | 0 (0.0) |
| Farmer/Agriculture | 1 (3.3) | 0 (0.0) | 1 (10.0) | 0 (0.0) |
| Freelancer | 3 (10.0) | 0 (0.0) | 0 (0.0) | 3 (30.0) |
| Public Company Employee/State Enterprise | 4 (13.3) | 0 (0.0) | 2 (20.0) | 2 (20.0) |
| Soldier | 1 (3.3) | 1 (10.0) | 0 (0.0) | 0 (0.0) |
| Student | 2 (6.7) | 0 (0.0) | 2 (20.0) | 0 (0.0) |
| Unemployed | 14 (46.7) | 9 (90.0) | 4 (40.0) | 1 (10.0) |
| Housing | | | | |
| House | 20 (66.7) | 10 (100.0) | 10 (100.0) | 0 (0.0) |
| Apartment | 10 (33.3) | 0 (0.0) | 0 (0.0) | 10 (100.0) |
| Socioeconomic Class | | | | |
| Lower | 5 (16.7) | 3 (30.0) | 0 (0.0) | 2 (20.0) |
| Middle | 25 (83.3) | 7 (70.0) | 10 (100.0) | 8 (80.0) |
| Num. Children in Household | | | | |
| 1-2 | 13 (43.3) | 6 (60.0) | 4 (40.0) | 3 (30.0) |
| 3-4 | 13 (43.3) | 3 (30.0) | 5 (50.0) | 5 (50.0) |
| 5+ | 4 (13.3) | 1 (10.0) | 1 (10.0) | 2 (20.0) |

| | | | | |
|---|---|---|---|---|
| Num. Domestic Water Containers in Home | | | | |
| 1-2 | 22 (73.3) | 7 (70.0) | 8 (80.0) | 7 (70.0) |
| 3-4 | 7 (23.3) | 3 (30.0) | 2 (20.0) | 2 (20.0) |
| 5+ | 1 (3.3) | 0 (0.0) | 0 (0.0) | 1 (10.0) |
| Frequency of Bednet Use | | | | |
| Never | 9 (30.0) | 3 (30.0) | 2 (20.0) | 4 (40.0) |
| Rarely | 0 (0.0) | 0 (0.0) | 0 (0.0) | 0 (0.0) |
| Regularly/Most of the Time | 4 (13.3) | 0 (0.0) | 0 (0.0) | 4 (40.0) |
| All of the time | 17 (56.7) | 7 (70.0) | 8 (80.0) | 2 (20.0) |
| Larvicide Use in Home | | | | |
| No | 28 (93.3) | 10 (100.0) | 10 (100.0) | 8 (80.0) |
| Yes | 2 (6.6) | 0 (0.0) | 0 (0.0) | 2 (20.0) |
| Insecticide Use in Home | | | | |
| No | 19 (63.3) | 6 (60.0) | 5 (50.0) | 8 (80.0) |
| Yes | 11 (36.7) | 4 (40.0) | 5 (50.0) | 2 (20.0) |
| Mosquito Coil Use in Home | | | | |
| No | 10 (33.3) | 7 (70.0) | 2 (20.0) | 1 (10.0) |
| Yes | 20 (66.7) | 3 (30.0) | 8 (80.0) | 9 (90.0) |
| Ever Had Dengue Infection | | | | |
| No | 27 (90.0) | 9 (90.0) | 10 (100.0) | 8 (80.0) |
| Yes | 3 (10.0) | 1 (10.0) | 0 (0.0) | 2 (20.0) |
| Insect Bites in Last 30 Days | | | | |
| No | 2 (6.7) | 0 (0.0) | 1 (10.0) | 1 (10.0) |
| Yes | 28 (93.3) | 10 (100.0) | 9 (90.0) | 9 (90.0) |
| Mean Num. Bites (Min, Max) | 5.9 (3, 10) | 4.9 (3, 7) | 7.1 (5, 10) | 5.6 (3, 8) |
| Redness | | | | |

| | | | | |
|---|---|---|---|---|
| No | 7 (23.3) | 0 (0.0) | 6 (60.0) | 1 (10.0) |
| Yes | 23 (76.7) | 10 (100.0) | 4 (40.0) | 9 (90.0) |
| Swelling | | | | |
| No | 1 (3.3) | 0 (0.0) | 1 (10.0) | 0 (0.0) |
| Yes | 29 (96.7) | 10 (100.0) | 9 (90.0) | 10 (100.0) |
| Mean Bite Size, mm (S.D.) | | | | |
| 15 min | 5.0 (2.8) | 4.4 (1.2) | 6.2 (4.1) | 4.4 (2.3) |
| 30 min | 4.8 (3.3) | 4.1 (1.3) | 6.0 (4.8) | 4.3 (2.7) |
| 4 hours | 2.2 (1.2) | 1.9 (1.1) | 2.6 (0.9) | 2.0 (1.5) |
| 48 hours | 1.1 (1.2) | 1.6 (1.4) | 1.3 (1.1) | 0.5 (0.7) |
| Mean OD (S.D.) to *Aedes aegypti* SGE | | | | |
| Day 0 | 0.16 (0.08) | 0.15 (0.06) | 0.17 (0.10) | 0.16 (0.07) |
| Day 14 | 0.17 (0.07) | 0.15 (0.06) | 0.17 (0.10) | 0.17 (0.06) |
| Positive on PanBio® Dengue Indirect IgG | 30 (100) | 10 (100) | 10 (100) | 10 (100) |

Table S1. **Cohort Demographics by Sample Evaluation Modality**. All data presented as n (%) unless otherwise stated. OD = optical density SGE = salivary gland extract.

Table S2**. List of differentially expressed genes in excel format.**

Table S3. **List of over represented pathways at the Reactome database in excel format**.

Table S4. **DEG under an FDR cutoff <0.05 and log2FC >1 in at least one timepoint in excel format**

| Fluorochrome | Innate 1 | Innate 2 | Adaptive |
|---|---|---|---|
| **BUV395** | CD3 / Clone SK7 | | CD3 / Clone SK7 |
| **BUV496** | CD4 / Clone SK3 | | CD4 / Clone SK3 |
| **BUV737** | | CD16 / Clone 3G8 | CCR7 / Clone 2-L1-A |
| **BV421** | | CD163 / Clone GHI/61 | CD45RA / Clone HI100 |
| **BV605** | | CD14 / Clone 63D3 | CLA / Clone HECA-452 |
| **BV650** | CD1c / Clone L161 | | CCR4 / Clone 1G1 |
| **BV711** | | CD56 / Clone 5.1H11 | CXCR3 / Clone G025H7 |
| **BV785** | CD69 / FN50 | CD69 / FN50 | CD69 / FN50 |
| **BB515** | | CD117 / Clone 104D2 | |
| **PE-Texas Red** | | | CD45 |
| **PE** | CD207 / Clone 10E2 | | |
| **BB700** | | | |
| **PE-Cy7** | CD45 / HI30 | CD45 / HI30 | PD1 / Clone EH12.1 |
| **APC** | | | |
| **APC-R700** | CD25 / Clone 2A3 | CD25 / Clone 2A3 | CD25 / Clone 2A3 |
| **APC-H7** | CD8 / Clone SK1 | | CD8 / Clone SK1 |
| **Amcyan** | **Viability** | **Viability** | **Viability** |

Table S5**. List of antibodies used for flow cytometry analysis, clones and fluorochromes.**

**<u>Supplementary Figures</u>**

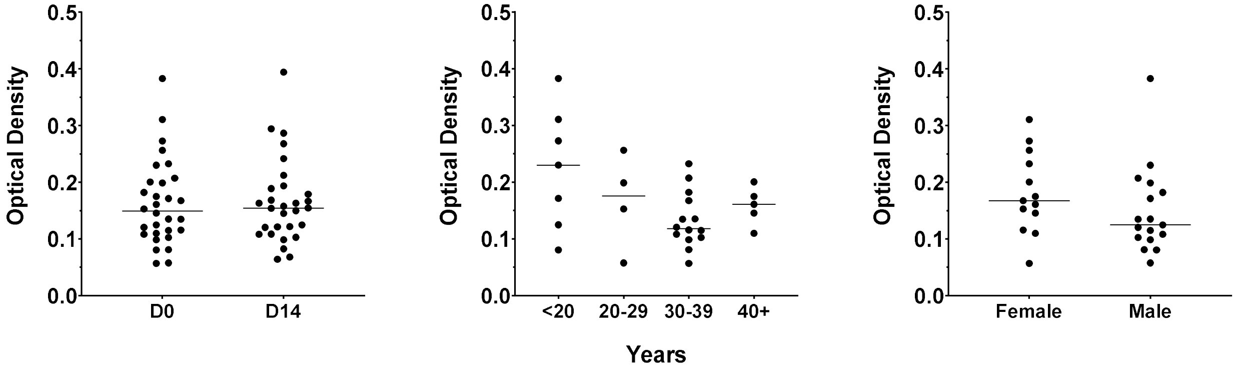

Figure S1. **Change in optical density of anti-SGE IgG by time, age, and gender as measured by Aedes aegypti salivary IgG ELISA.** No significant differences were noted in the groups. Line represents median

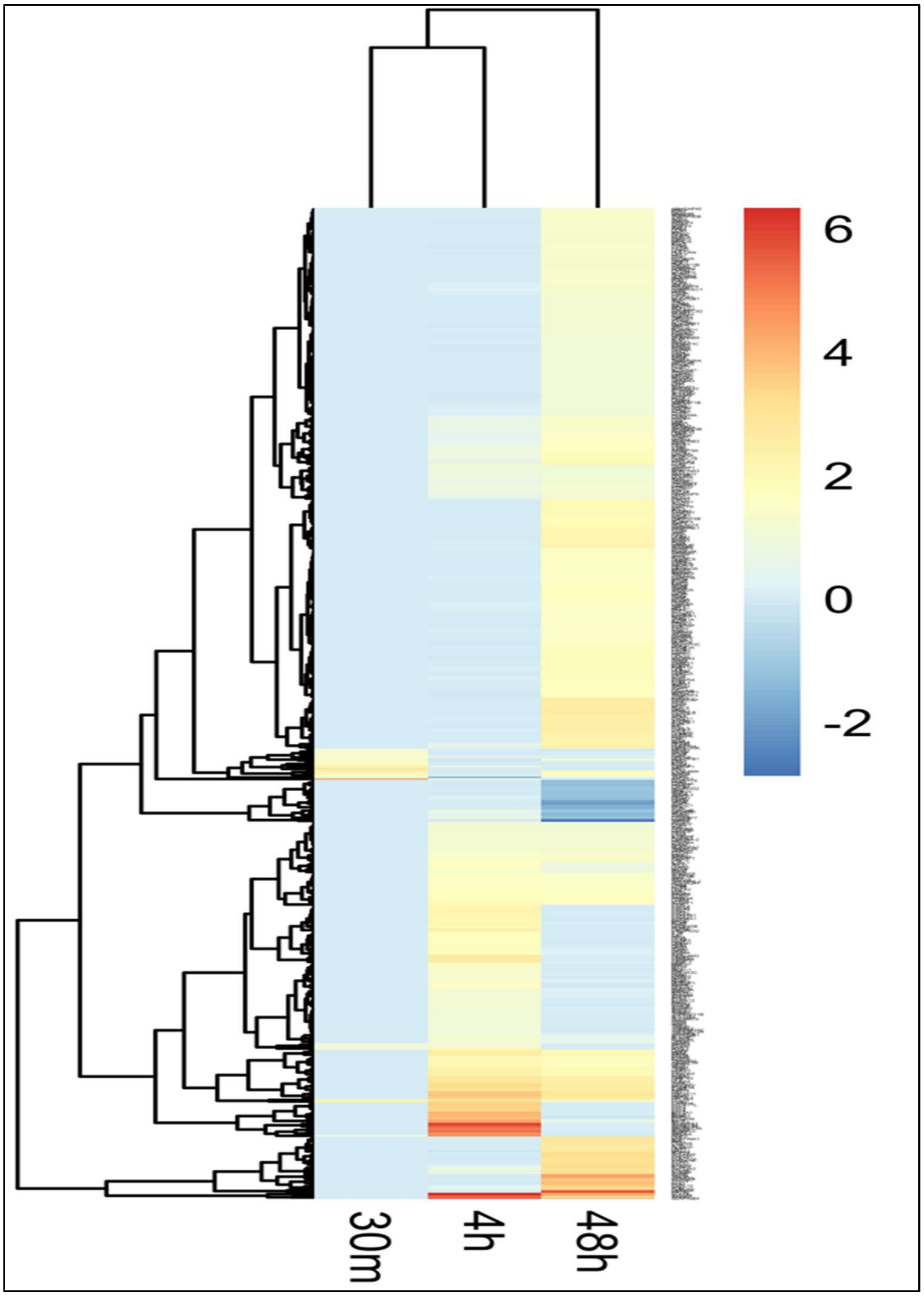

Figure S2**. Comprehensive heat map of all differentially expressed genes at all time points**.

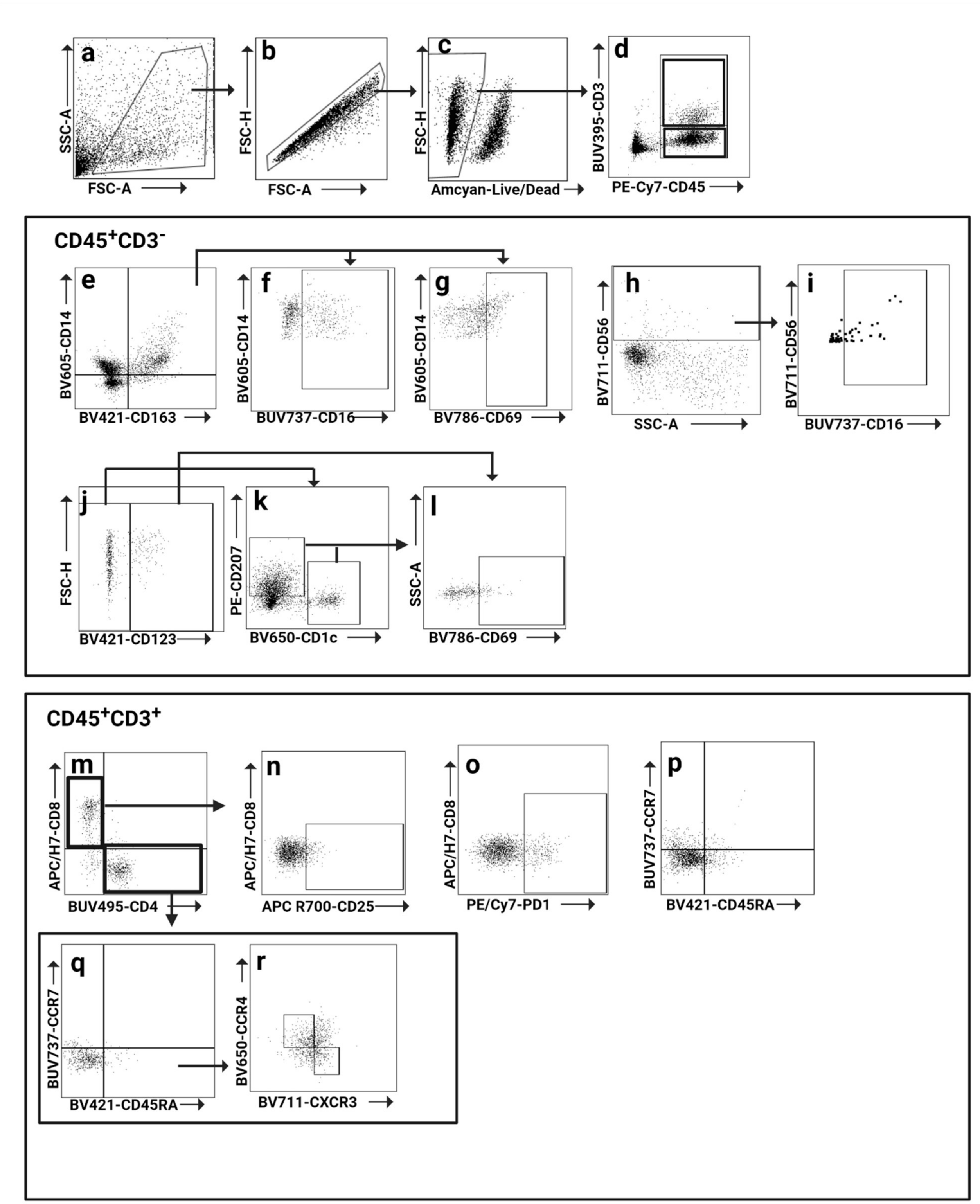

Figure S3. **Flow cytometry gating strategy.** **(a)** All cells **(b)** Singlets **(c)** Live cells **(d)** $CD3^+$ and $CD45^+$ gates **(e)** M2MØ gate ($CD14^+CD163^+$) **(f)** M2MØ $CD16^+$ **(g)** Activated M2MØ **(h)** NK cells **(i)** NK cells $CD16^+$ **(j)** Plasmacytoid DCs **(k)** Langerhans cells ($CD207^+$) dermal DCs ($CD1c^+$) **(l)** Activated Langerhans cells/dermal DCs **(m)** $CD8^+$ and $CD4^+$ T cells gates **(n)**

Activated CD8$^+$ T cells **(o)** PD1$^+$ CD8$^+$ T cells **(p- q)** Central memory (CCR7$^+$CD45RA$^-$), Naïve memory (CCR7$^+$CD45RA$^+$), Effector memory (CCR7$^-$CD45RA$^-$) and Terminal differentiated effector memory (CCR7$^-$CD45RA$^+$) **(r)** Th17/Th2 (CCR4$^+$CXCR3$^-$) and Th17/Th1 (CCR4$^-$CXCR3$^+$) compartments.

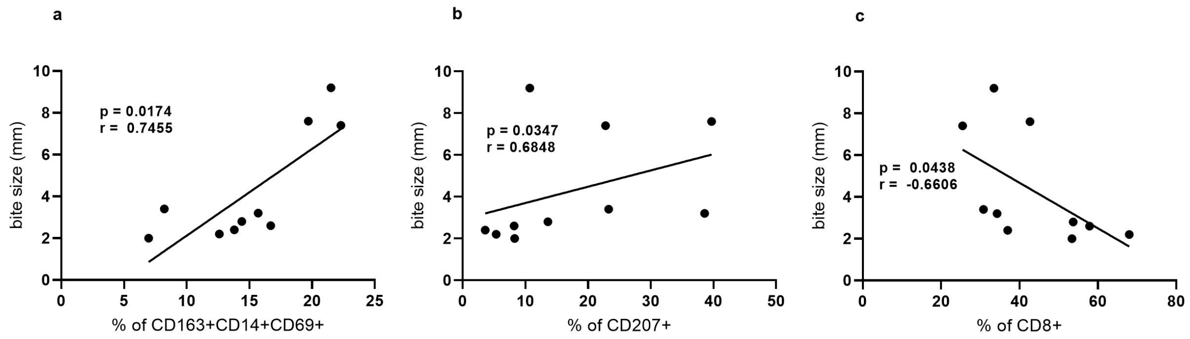

Figure S4. **Clinically observed changes in 'bite size' correlate to various immunological changes. (a)** Positive correlation between bite size at 30 minutes and frequency of activated M2 macrophages (CD163+CD14+CD69+) and Langerhans cells **(b)**. **(c)** Negative correlation between bite size at 30 minutes and frequency of CD8+ T cells at 48 hours**.** Statistical analysis were performed with Pearson correlation coefficient.

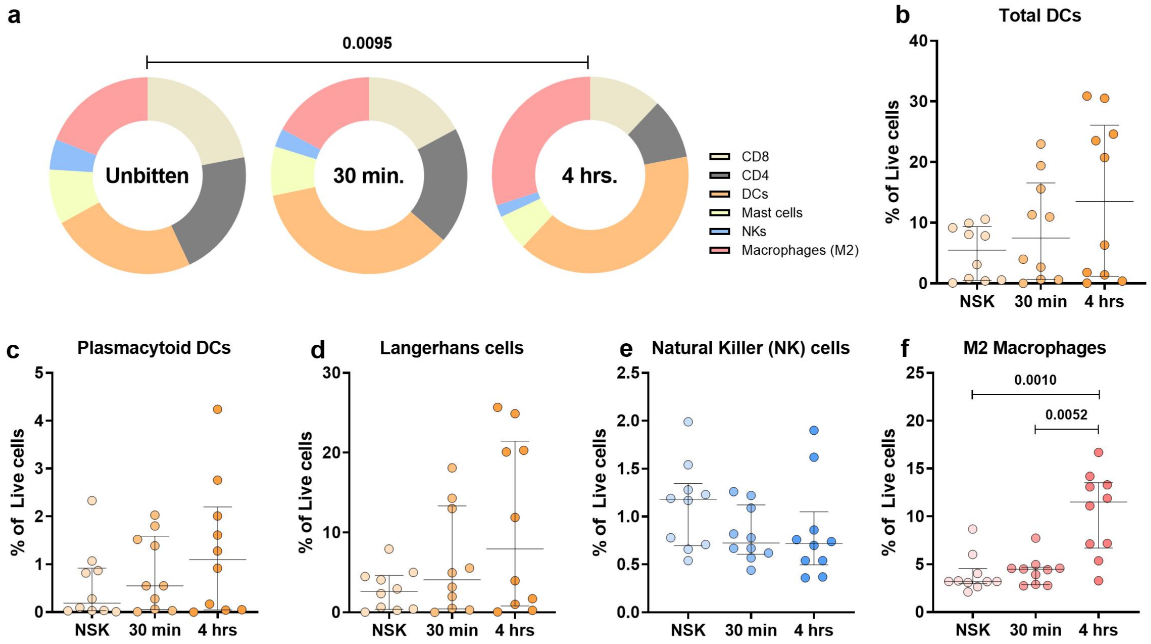

Figure S5. **a,** Pie charts showing changes in the frequencies of skin immune cells during early innate immune response to mosquito bite at 30 minutes and 4 hours after exposure. **b-f**

Frequencies are reported as percentages of total live cells. Statistical analysis were performed with Chi-square test (**a**) and Friedman + Dunn's multiple comparisons test (**b-f**). Bars indicate median and interquartile range.

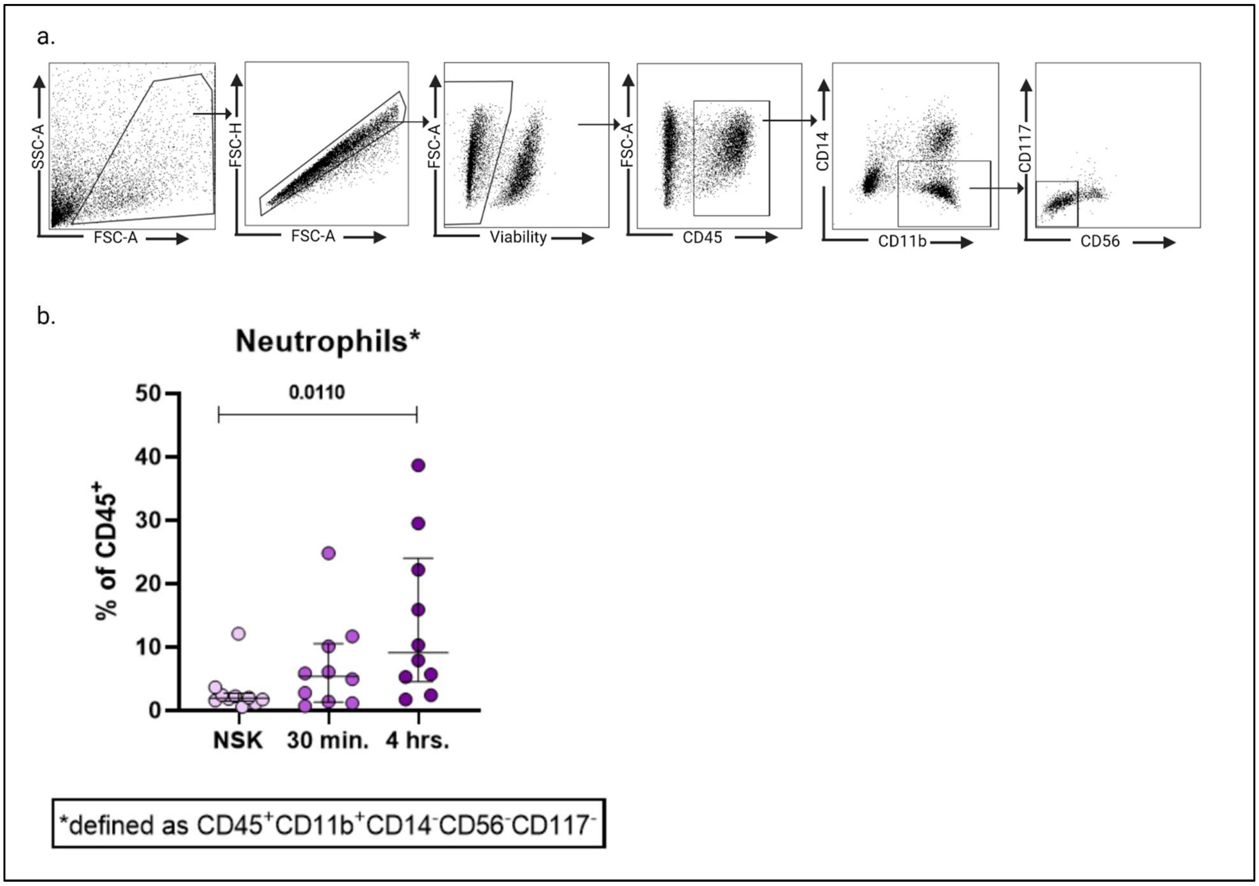

Figure S6. **Changes in myeloid cells CD45+CD11b+CD14-CD56-CD117- frequencies. (a)** Gating strategy defining the population of myeloid cells excluding lymphocytes, monocytes/macrophages, NK cells and mast cells. **(b)** Change in the frequency of neutrophils – defined as CD45+CD11b+CD14-CD56-CD117-. Statistical analysis were performed with Wilcoxon signed-rank test. Bars indicate median.

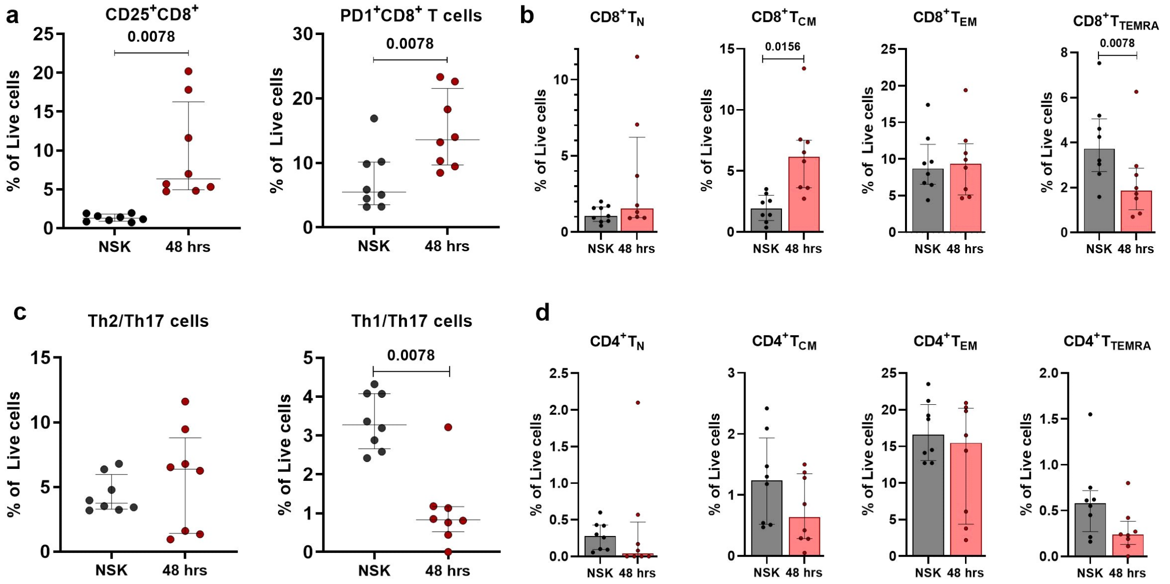

Figure S7. Changes in frequencies of CD4+ and CD8+ T cell populations as percentages of total live cells. Statistical analysis was performed with Wilcoxon signed-rank test. Bars indicate median and interquartile range.

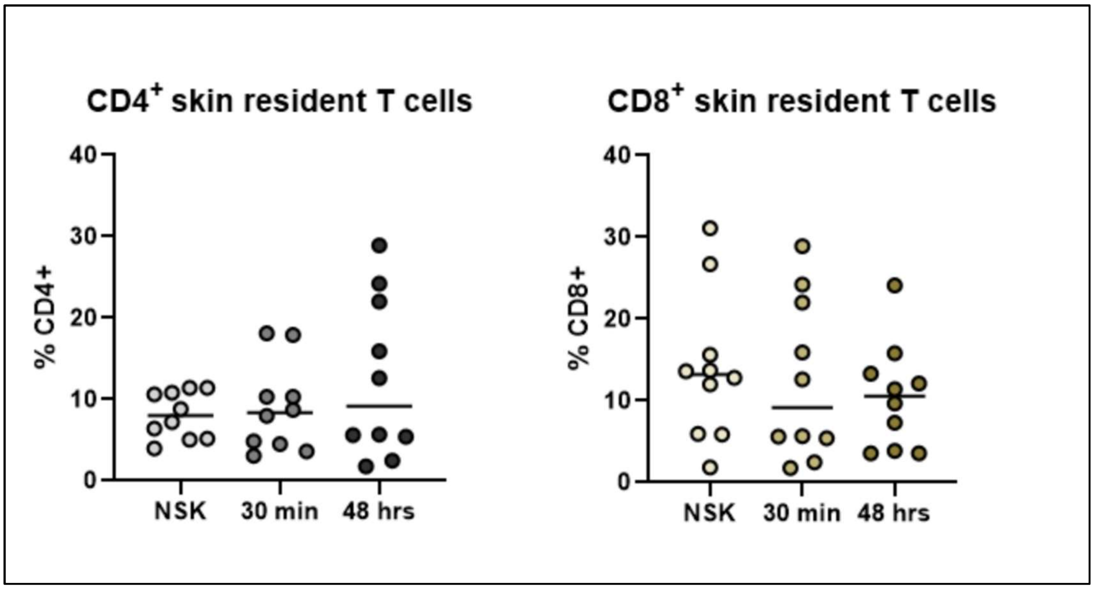

Figure S8. **No changes are observed in the frequencies of resident T cells at any time point post mosquito bite.** **(a)** Frequencies of CD4+CD103+CLA+ and **(b)** CD8+CD103+CLA+ T cells in normal skin and from biopsies at 30 minutes and 4 hours post-bite. Statistical analysis were performed with Wilcoxon signed-rank test. Bars indicate median.

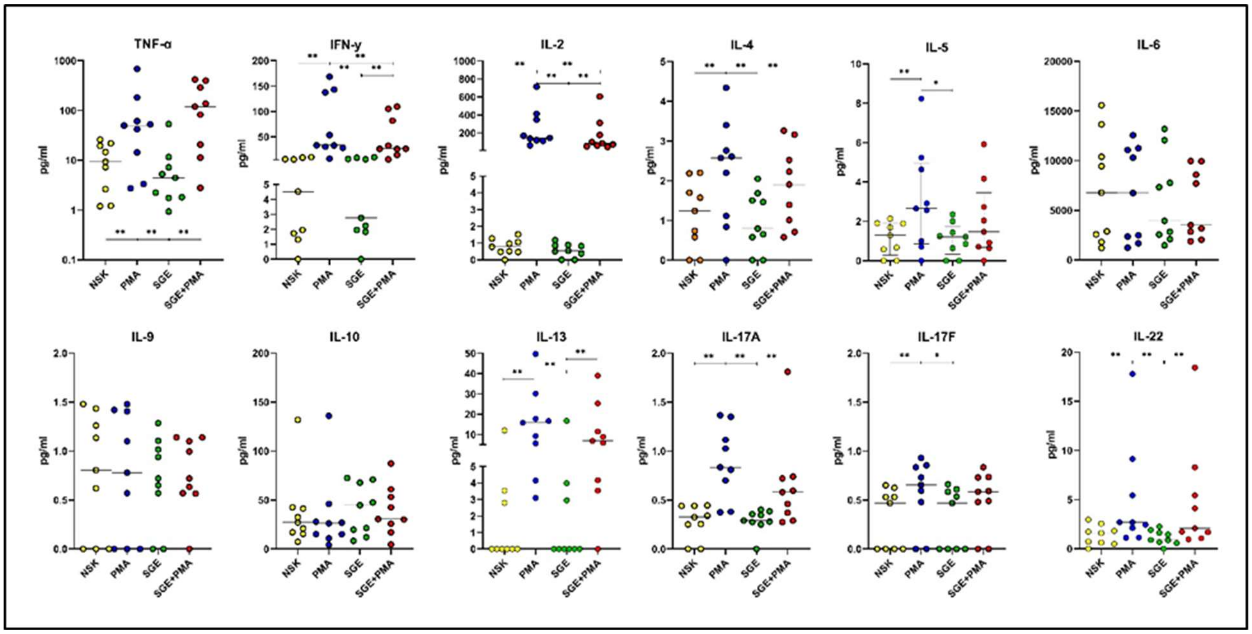

Figure S9. **Cytokine production after stimulation with PMA/Ionomycin preparation.**

Cytokine production measured in skin cell culture supernatant after treatment with PMA/Ionomycin in the presence or absence of SGE. Skin dissociated cells were seeded on round bottom 96-well plates (50.000 cells/well) and treated with SGE (10 µg/mL) or PBS for 24 hours. Cells were stimulated with PMA (0.1µg/mL) and Ionomycin (1µg/mL) or left unstimulated for the last 6 hours of the culture. Statistical analysis was performed with Wilcoxon signed-rank test. N=9 individuals. Bars indicate median. *$p<0.05$, **$p<0.01$, ***$p<0.001$, ****$p<0.0001$
